# Different spatial profiles of aberrant N-glycans in pediatric and adult MOGHE brain tissue

**DOI:** 10.64898/2026.08.12.744424

**Authors:** Chiara Calabretta, Dalia De Santis, Grace Grimsley, Greta De Cicco, Laura Rossini, Margherita Marchi, Ilaria D’Amato, Ester Cifaldi, Michele Rizzi, Gianluca Marucci, Laura Tassi, Francesco Cardinale, Francesca Ragona, Roberta Di Giacomo, Niccolò D’Agaro, Giulia Capitoli, Marco de Curtis, Richard R. Drake, Rita Garbelli, Cinzia Cagnoli

## Abstract

Mild malformation of cortical development with oligodendroglial hyperplasia in epilepsy (MOGHE) is a recently recognized epilepsy-associated lesion frequently linked to brain-restricted somatic variants in *SLC35A2*, a gene encoding the Golgi UDP-galactose transporter. Although previous studies demonstrated altered glycosylation in *SLC35A2*-mutated MOGHE tissue, the spatial relationship between glycosylation defects and histopathological abnormalities remains poorly understood. We applied matrix-assisted laser desorption/ionization mass spectrometry imaging (MALDI-MSI) using formalin-fixed paraffin-embedded brain tissue from six histologically confirmed MOGHE cases (three pediatric and three adult) and three temporal lobe epilepsy with hippocampal sclerosis (TLE-HS). We spatially evaluated N-glycan profiles across diagnostic tissue groups, with particular attention to molecular differences between lesional and perilesional regions and to recurrent abundance trends. All MOGHE cases harboured somatic *SLC35A2* variants. Histologically, oligodendroglial hyperplasia and heterotopic neurons were present in all cases, while patchy hypomyelination was restricted to pediatric cases.

Unsupervised spatial segmentation, integrated with neuropathological evaluation, revealed marked molecular heterogeneity in pediatric MOGHE. In these cases, lesional and perilesional regions were clearly distinguishable in both white matter (WM) and overlying grey matter (GM) boundaries patterns, whereas adult MOGHE and TLE-HS mainly showed a clearcut separation between WM and GM.

Spatial analysis confirmed enrichment of the previously reported aberrant N-glycan species *m/z* 2094 and, to a lesser extent, *m/z* 2297 within MOGHE tissue, particularly in pediatric lesional WM. Notably, the distribution of *m/z* 2094 closely overlapped with areas of hypomyelination. Quantitative trajectory analysis of 151 detected N-glycan ions identified recurrent abundance profiles. Three representative spatial patterns emerged: pediatric lesion-enriched, pediatric perilesion-enriched, and TLE-HS-enriched profiles. Pediatric lesions were characterized by increased abundance of multiantennary glycans lacking terminal galactose residues and reduced abundance of galactosylated biantennary and multiantennary structures, consistent with defective UDP-galactose transport. In contrast, adult lesional and perilesional tissues exhibited largely overlapping glycomic profiles.

These findings provide the first spatially resolved evidence that glycosylation abnormalities in *SLC35A2*-mutated MOGHE are closely associated with lesional pathology, particularly hypomyelination, and are substantially more pronounced in pediatric than adult cases. Spatial glycomics may therefore offer new insights into MOGHE pathophysiology and support the development of targeted therapeutic approaches aimed at correcting galactosylation defects.

## Introduction

Epilepsy surgery represents an effective treatment option in patients with focal drug-resistant epilepsy (DRE)^1^. The spectrum of structural brain lesions amenable for surgery includes malformations of cortical development (MCD), among which focal cortical dysplasias (FCD) are the most common entities, particularly in the pediatric population.^2^ FCDs comprise a heterogeneous group of brain malformations, some of which are genetically determined, caused by abnormalities in neuronal migration and cortical organization. In the updated FCD classification of the International League Against Epilepsy (ILAE) the mild malformation of cortical development with oligodendroglial hyperplasia in epilepsy (MOGHE) was formally recognized as a new pathological entity.^3^ Histopathologically, MOGHE is characterized by *i)* increased oligodendroglial density at the grey-white matter junction, *ii)* patchy hypomyelination, and *iii)* the presence of heterotopic neurons within the white matter. The disorder predominantly involves the frontal lobe (approximately 50–70% of cases), although temporal and multilobar involvement has also been reported. Surgical series indicate that about half of patients achieve seizure freedom after resection,^4^ underscoring the need for a better understanding of disease mechanisms and for personalized treatment.

Recent studies have identified brain-restricted somatic variants in *SLC35A2* in approximately half of histologically confirmed MOGHE cases. *SLC35A2*, located on chromosome Xp11.23, encodes a uridine diphosphate (UDP)-galactose transporter localized in the Golgi apparatus. Pathogenic variants impair UDP-galactose uptake into Golgi vescicles, leading to both defective protein glycosylation and the formation of truncated N-glycan structures in affected brain tissue.^5^ Glycosylation is an essential biological process and a common post-translational modification of proteins and lipids. Both N- and O-linked protein glycosylation are key contributors to the cellular glycome; notably, N-glycosylation occurs in more than half of all proteins and is critical for proper protein folding, localization, stability and activity.^6^ A direct link between altered glycosylation pattern and epilepsy is supported by genetic disorders of glycosylation (CDG) associated with germline *SLC35A2* variants, that are clinically characterized by a broad neurological phenotype, featuring intellectual disability, hypotonia, epileptic spasms and DRE.^7^ In this context, galactose supplementation has been proposed as a rational therapeutic strategy for individuals with *SLC35A2*-CDG.^8^

In MOGHE cases, aberrant N-glycan profiles in *SLC35A2*-mutated brain samples have been demonstrated using nanoLC-MS/MS-based glycomics^9^; more recent glycoproteomic and chemo-enzymatic histological studies confirmed glycosylation defects in these tissues.^10^ Despite these advances, no studies have yet directly linked altered glycosylation patterns to define MOGHE histopathological features, particularly white matter abnormalities and oligodendroglial hyperplasia.

In this context, matrix-assisted laser desorption/ionization mass spectrometry imaging (MALDI-MSI) has emerged as a powerful analytical platform for spatially resolved N-glycan profiling directly within tissue sections. N-glycan MALDI-MSI enables the generation of high-resolution molecular maps linking specific glycosylation patterns to distinct anatomical regions and pathological features, including brain,^11,12^ providing unprecedented insight into disease-specific mechanisms.^13^

On these premises, we performed a spatially resolved glycomic analysis on formalin fixed and paraffin embedded (FFPE) brain tissue sections from pediatric and adult MOGHE cases, compared to temporal lobe epilepsy with hippocampal sclerosis (TLE-HS) cases as control, to investigate where glycosylation defects arise and how they relate to histopathological abnormalities. We evaluated N-glycan profiles across diagnostic tissue groups, with particular attention to molecular differences between lesional and perilesional regions and to recurrent abundance trends in pediatric and adult MOGHE. These findings may contribute to a better understanding of MOGHE pathophysiology and provide a framework for the future studies of targeted and personalized therapeutic strategies.

## Materials and methods

### Patient’s selection and tissue characterization

We retrospectively considered nine patients who underwent surgery for DRE in frontal and temporal lobe at the *Carlo Besta* Neurological Institute Foundation and at the *Claudio Munari* Epilepsy Surgery Center at Niguarda Hospital (both in Milan, Italy) following comprehensive electroclinical and magnetic resonance imaging (MRI) evaluation. The ethics committees of both institutions approved the diagnostic and therapeutic procedures and the use of brain material for research (EpiBesta protocol n.83, 14-04-2021; Niguarda protocol n. 696-16112021). Histopathological examination of FFPE frontal and temporal lobe tissue sections was performed in six cases (three pediatric and three adult) that met the criteria for MOGHE diagnosis according to the latest FCD classification.^3^ Temporal neocortex tissue from three additional adult cases of TLE-HS was included as control.

All specimens were processed using standard fixation and inclusion protocols. Serial sections were processed for histology (hematoxylin and eosin-H&E, cresyl violet and Luxol fast blue staining-LFB) and immunohistochemistry was performed according to the ILAE workup guidelines for epilepsy surgery specimens.^14^ In MOGHE cases, tissue levels were selected to include areas exhibiting both pronounced histopathological alterations (lesional core) and perilesional tissue, either within the same section or in different tissue blocks obtained from the same patient. Perilesional tissue was utilized as same-patient internal control against the lesional area in subsequent glycan analysis. The density of Olig2-positive cells was manually quantified within three predefined white matter (WM) Region of Interests (ROIs) of similar size (0.09 mm^2^) in both lesional core and perilesional area in MOGHE cases, as well as in the additional temporal neocortex control tissue from TLE-HS, using ImagePro Premier Software (v.9.3, Media Cybernetics, Rockville, MD).

For MOGHE cases, histological diagnoses were further integrated with genetic findings. Genomic DNA was extracted from FFPE brain tissue sections enriched for lesional core in five out of six cases, using the QIAamp DNA FFPE Advanced UNG Kit (QIAGEN S.r.l.) according to manufacturer’s instructions. DNA quality was assessed prior to library construction to ensure an appropriate fragment size; in this context, the DNA Integrity Number (DIN) was evaluated using a size distribution analyser. Where necessary, DNA fragmentation was performed using a Covaris® shearing instrument to obtain fragments with an average insert size of 150–300 bp. Subsequently, 500 ng of fragmented DNA was analysed using a custom targeted next-generation sequencing (NGS) approach, including 13 candidate genes panel (xGen™ MRD Hyb Panel, IDT) for FCD (Supplementary Table 1). DNA quantity was measured using QUBIT (ThermoFisher), while integrity and quality were evaluated using Genomic DNA D1000 ScreenTape Analysis, in conjunction with the TapeStation systems (Agilent). Libraries were prepared using the xGen cfDNA & FFPE DNA Library Prep v2 MC (IDT), enriched with xGen™ Hybridization Capture Core Reagents (IDT) and sequenced on a NextSeq550 platform (Illumina). In the remaining case, fresh frozen tissue was analysed with a target FCD panel (Supplementary Table 1) using a single molecule Molecular Inversion Probes (smMIPs).^15^

### N-glycan sample preparation for MALDI-MSI

For N-glycan analysis, 7-μm-thick sections were mounted on indium tin oxide slides. FFPE sections were dewaxed at 60° C for 1 h, followed by a series of washes with xylene, ethanol, and HPLC-grade water to deparaffinize and rehydrate the tissue. Antigen retrieval using citraconic acid (pH 3.0) for 30 min enhanced peptide N-glycosidase F (PNGaseF PRIME) enzyme accessibility. Slides were subsequently coated with 0.1 μg/μl PNGaseF (N-zyme Scientific, Doylestown, PA, USA) using a TM Sprayer (HTX Technologies, Chapel Hill, NC, USA).^16^ A 7 mg/ml MALDI matrix solution of α-cyano-4-hydroxycinnamic acid in 50% acetonitrile/0.1% trifluoroacetic acid was applied to the de-glycosylated tissues (20 passes of matrix at 50 µl/min flow rate). Following MSI acquisition, the matrix was removed, and the same tissue sections were stained with H&E.

### MALDI-MSI parameters and data processing

Released N-glycans were analysed over a mass range of 700-4000 *m/z* in positive-ion mode using a timsTOF fleX MALDI-QTOF mass spectrometer (Bruker Corporation, Billerica, MA, USA). Data acquisition was performed with a laser spot size of 20 μm, 300 laser shots per pixel, and a raster size of 40 μm. Spectral processing and analysis were carried out using SCiLS Lab software (v.2026aPro, Bruker Corporation). N-glycan spectra were converted to relative intensities by normalizing individual peak intensities to the Total Ion Count (TIC) for each sample. Signal-to-noise (S/N) ratios were evaluated by comparison with neighbouring *m/z* signals and matrix-related peaks; no fixed S/N threshold was applied.

N-glycan peaks present in the whole dataset were annotated by comparing *m/z* species to an in-house curated list composed of 176 *m/z*. This list was generated integrating brain N-linked glycans reported in human studies^11,17,18^ and in papers specifically addressed to MOGHE pathology.^9,10,19^ Since an MS/MS-based structural characterization was not performed, all assignments were reported as putative structures.

### Statistical analysis of MALDI-MSI data

Spatial segmentation was performed using SCiLS Lab software to identify tissue regions with distinct N-glycan profiles. The resulting segmentation maps were independently reviewed by two experienced neuropathologists to assess their concordance with histopathological features, including oligodendroglial and myelin alterations. ROIs were manually defined in SCiLS Lab according to histological features and spatial segmentation results and showed in Fig. 3E, H, M. Ion intensities were extracted from the selected ROIs and log₂(x+1) transformed before statistical analysis.

For trajectory analysis, N-glycan abundance was summarized across five tissue groups: TLE-HS tissue, adult MOGHE perilesional tissue, adult MOGHE lesional core tissue, pediatric MOGHE perilesional tissue, and pediatric MOGHE lesional core tissue. Grey- and white-matter measurements belonging to the same patient and region were averaged to obtain patient-level abundance profiles. For each *m/z* species, mean log₂-transformed intensities were standardized within feature to describe the relative abundance trajectory across the five groups. Grey and white matter (GM, WM) were retained separately for heatmap visualization. Before trajectory analysis, original MALDI-MSI signals were evaluated according to mean abundance, abundance range across the five tissue groups, and detection rate. Ions with *m/z* ≥3000 were excluded because this mass range was predominantly represented by signals within the lowest quartile of both mean abundance and between-group range. This step limited the influence of low-intensity signals whose relative variation could be amplified by within-feature standardization.

Standardized five-group trajectories were grouped by k-means clustering to identify recurrent abundance profiles. Differences among TLE-HS, adult MOGHE, and pediatric MOGHE were evaluated using the Kruskal-Wallis test, with p-values adjusted using the Benjamini–Hochberg false discovery rate. The magnitude of the group effect was quantified using epsilon squared (ε²).

Given the exploratory design and limited sample size, the final selection was not based on statistical significance alone. Candidate ions were evaluated by jointly considering ε², original signal abundance, five-group abundance range, detection rate, and consistency between their quantitative trajectory and spatial tissue distribution. Fifteen ions were selected as representative of three profiles: pediatric lesion-enriched, pediatric perilesion-enriched, and TLE-HS-enriched. The signals *m/z* 2094 and *m/z* 2297, previously described as aberrant in MOGHE tissue by Sim et al.^9^, were retained for comparison purpose. Signal-quality summaries, trajectory-cluster characteristics, and the quantitative metrics of the 15 representative ions are reported in Supplementary Table 2.

For selected *m/z* species, the relative intensity measured in lesional WM was correlated with the *SLC35A2* variant allele frequency (VAF) using the Spearman’s rank correlation coefficient.

All statistical analyses were performed with SCiLs Lab (Bruker), Prism (GraphPad Software, www.graphpad.com), and open-source R software 4.2.2 (R Foundation for Statistical Computing).

## Results

### Clinical and pathological differences among diagnostic groups

In pediatric MOGHE cases, the mean age at seizure onset was 1 year, the mean age at surgery was 4 ± 1.4 years and the mean epilepsy duration was 3 ± 1 years; in adult MOGHE, the mean seizure onset was 9 ± 4.2 years, the mean age at surgery was 44.5 ± 3.5 years and the mean epilepsy duration was 30.7 ± 10 years. Adult patients with TLE-HS, selected for comparative purpose, showed a mean age at seizure onset of 9.3 ± 9.1 years, a mean age at surgery of 44.3 ± 13.1 years and the mean epilepsy duration was 35 ± 4.4 years. Postsurgical outcomes at follow-up longer than 12 months were Engel class IA in 2/3 pediatric MOGHE, class I in 2/3 adult MOGHE and class IA in 2/3 TLE-HS. Among the MOGHE cases, surgical resection involved the frontal lobe in three patients and the temporal lobe in the remaining three (Table 1).

**Table 1.** Summary of the main clinical and genetic data.

| **I**<br>**D** | **Histolog**<br>**y** | **Male/**<br>**Female** | **Age at**<br>**surgery**<br>**(y)** | **Age at**<br>**onset**<br>**(y)** | **Outcom**<br>**e Engel**<br>**class** | **Site of**<br>**surger**<br>**y** | **VAF** | **Mutatio**<br>**n type** | **Nucleotide and protein**<br>**changes** |
| --- | --- | --- | --- | --- | --- | --- | --- | --- | --- |
| 1 | MOGHE | M | 3 | 1 | IA | F | 24% | missense | c.827T>C (p.Leu276Pro) |
| 2 | MOGHE | M | 4 | 1 | II | F | 39.5<br>% | frameshif<br>t | c.436_439dupCAGC<br>(p.Leu147fs) |
| 3 | MOGHE | F | 5 | 1 | IA | T | 13% | nonsense | c.553C>T (p.Gln185*) |
| 4 | MOGHE | M | 23 | 2 | IC | F | 36% | missense | c.844G>A (p.Gly282Arg) |
| 5 | MOGHE | M | 47 | 6 | III | T | 0.5% | missense | c.769G>A (p.Ala257Thr) – low confidence |
| 6 | MOGHE | M | 42 | 12 | IB | T | 8.7% | nonsense | c.553C>T (p.Gln185*) |
| 7 | TLE-HS | F | 34 | 1 | IA | T | na | na | na |
| 8 | TLE-HS | F | 40 | 8 | IA | T | na | na | na |
| 9 | TLE-HS | M | 59 | 19 | II | T | na | na | na |

**Legend:** **ID:** identification number; **VAF:** variant allele frequency; **y:** years.

On histological examination, all MOGHE cases showed WM abnormalities characterized by both increased density of oligodendroglial cells (Fig. 1A, C, G) and the presence of heterotopic neurons (Fig. 1F), consistent with the established diagnostic criteria. In areas defined as lesional core, higher Olig-2 cellularity was found (Fig. 1C and G) in comparison to perilesional tissue (Fig. 1E). Averaged oligodendroglial cell densities measured in both lesional core and perilesional tissue of the six MOGHE cases and in the three TLE-HS cases are illustrated in Fig. 1I. Furthermore, patchy areas of hypomyelination were observed exclusively in pediatric cases within the WM lesional core (Fig. 1D compared with adult 1H), in agreement with previously published data.^20^ Temporal neocortex samples in TLE-HS cases did not show microscopic alterations (data not shown).

**Figure 1.**
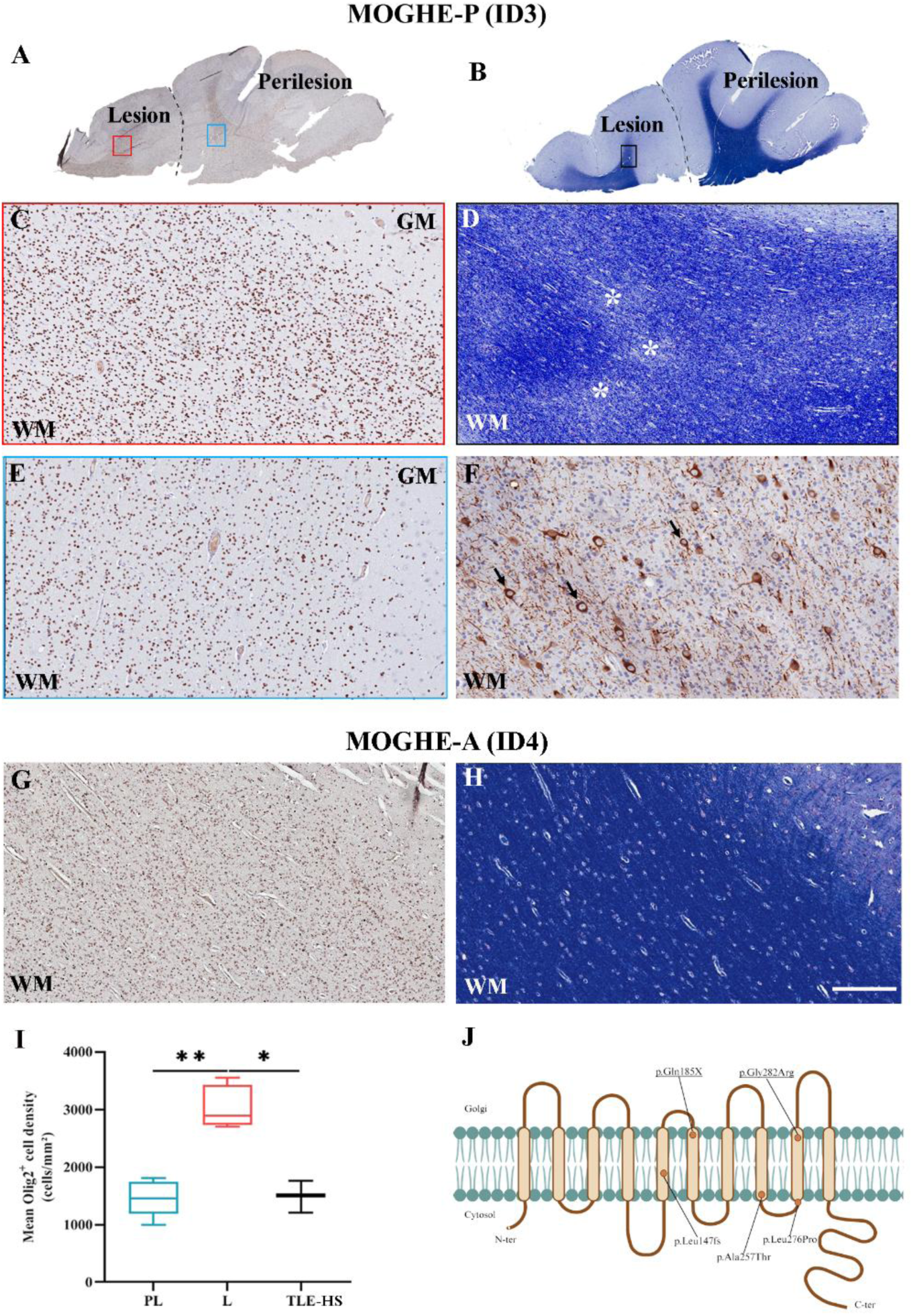
Histological features in pediatric and adult MOGHE cases. **A-F**: Representative tissue stainings from a pediatric MOGHE case demonstrating the characteristic histological hallmarks: increased density of Olig2-positive cells (**A**, **C**) and MAP2-positive heterotopic neurons (**F**), along with patchy areas of hypomyelination in the WM (**B**, **D**, LFB staining). The perilesional area (PL) included in the same section (**E**) was used for internal comparison. **G**, **H**: In an adult MOGHE case, Olig2-positive hyperplasia (**G**) is not associated with hypomyelination, as shown by LFB staining (**H**). **I**: Quantification in WM confirms that the lesional areas (L, in red) exhibit increased density of Olig2-positive cells across all MOGHE pediatric and adult cases, compared to perilesion (PL, in blue) and control TLE-HS samples (in black); L *vs* PL: ** p<0.001; L vs TLE-HS: * p<0.05. **J**: SLC35A2 protein showing the position of the somatic variants identified in our MOGHE cohort. Variants already described are underlined. Dashed lines in panels A and B indicate the histological boundaries between lesion and perilesion and boxes in panels A and B are shown at higher magnification in C, D and E, respectively. Case identifiers ID correspond to those reported in Table 1. Panel J created with BioRender. Scale bar: 5,9mm (A-B); 250 micron (C-E; G, H); 140 micron (F)

In all MOGHE cases, potentially pathogenic somatic mutations in the X-linked gene *SLC35A2* (NM_005660.3) were identified (Fig. 1J). The nonsense mutation p.Gln185* was found in two cases, one female with a VAF of 12% and in a male with VAF of 9%. One frameshift mutation p.Leu147fs was found in a male with a VAF of 39%, and three missense mutations (p.Leu276Pro, p.Gly282Arg, p.Ala257Thr) in three males, with 34.5, 24 and 0.5% of VAF, respectively. The extremely low VAF (0.5%) makes it challenging to establish whether the phenotype is attributable to such a small fraction of mutant cells. Table 1 summarised the main clinical and genetic information.

### Different glycosylation pattern across different diagnostic groups and anatomical regions

Unsupervised spatial segmentation was performed independently for each tissue section and interpreted together with the pathology-annotated H&E staining performed after MALDI-MSI and with immunohistochemical staining of adjacent sections.

In pediatric MOGHE, spectral segmentation distinguished not only WM from GM, but also lesional from perilesional regions in both WM and GM (Fig. 2A, B and Supplementary Fig. 1). In contrast, adult MOGHE, showed clear WM-GM separation, but no distinct segmentation between lesional and perilesional regions (Fig. 2C, D and Supplementary Fig. 1). TLE-HS samples highlighted only separation between WM and GM (Fig. 2E, F and Supplementary Fig. 1).

**Figure 2:**
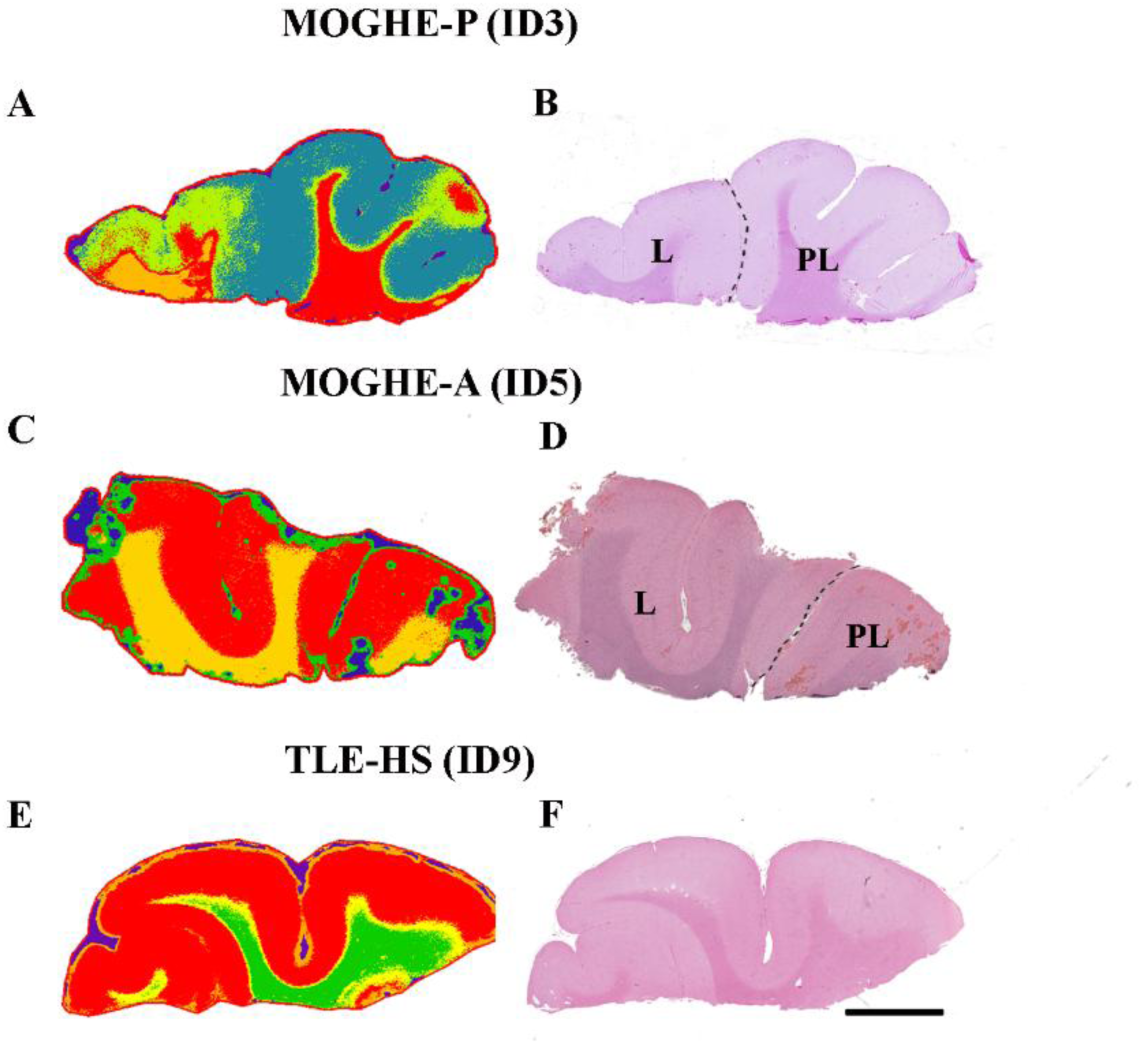
Unsupervised N-glycan segmentation analysis compared to adjacent H&E stained sections in different diagnostic groups. Bisecting k-means clustering applied to representative pediatric MOGHE (**A**), adult MOGHE (**C**) and control TLE-HS (**E**). Mass spectra are clustered based on similarity and visualized on tissue section as segmentation maps. In the pediatric case (**A**), the automatic spatial segmentation discriminates the WM from the GM and, notably, the WM lesional core (L, with Olig2 hyperplasia and hypomyelination at histology) versus the perilesion (PL); in GM also there is a differentiation between L and PL, despite any evident histological alteration. **B**: Post-MALDI Hematoxylin and Eosin (H&E) staining validates the segmentation map. **C**: In adult MOGHE, glycan analysis confirms a segregation between WM and GM, without a clear difference between L (presenting only WM Olig2 hyperplasia) and PL areas, visible on H&E staining (**D**). **E**: In control TLE-HS, only differentiation between WM and GW are obtained as validated in H&E (**F**). Dashed lines in panels C and E indicate the histological boundaries between L and PL. Case IDs correspond to those reported in Table 1. Scale bar: 4mm (B, D, F)

These distinct histology-integrated segmentation patterns guided ROIs definition for the subsequent quantitative analyses. Representative ROIs annotations are shown in Fig. 3E, H and M.

**Figure 3.**
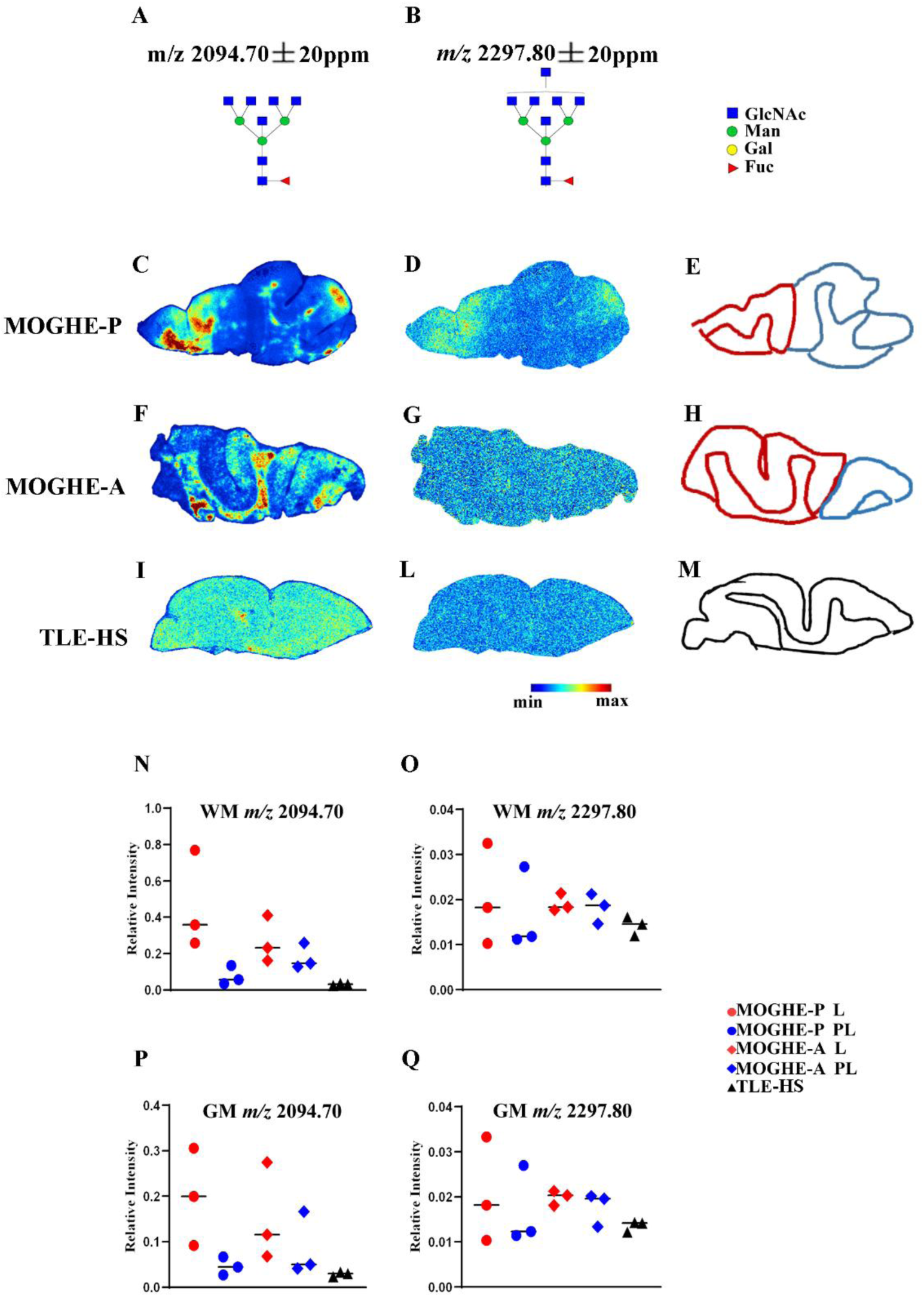
Spatial distribution and relative intensity of selected, literature-supported, N-glycans in different diagnostic groups. Molecular structures and corresponding legend of two selected *m/z* species previously reported as altered in brain tissue from MOGHE cases.^10^ The signal at *m/z* 2094 (Hex3HexNAc7Fuc1 in **A**) is markedly enriched in MOGHE cases (**C**, **F**) compared with control TLE-HS (**I**). Notably, the signal is predominantly localized within the WM and, in the pediatric case (**C**), is further enriched in the lesional area (indicated as a red area in **E, H**) relative to the adjacent, less affected perilesional area (blue area), as defined by histopathological evaluation. In contrast, *m/z* 2297 (Hex3HexNAc8Fuc1, in **B**) is detected at low levels across all cases, particularly in TLE-HS (**D**, **G**, **L**). **E, H, M**: Representative regions of interest (ROIs) used for quantitative analysis and annotated based on histological guidance and segmentation results. Specifically: red outlines indicate L areas, blue outlines indicate PL in MOGHE cases, and black outlines indicate GW and WM ROIs in control TLE-HS. **N-Q**: Dot plots showing the relative intensity quantifications of the two *m/z* species across the different ROIs and corresponding legend. Each dot represents the mean intensity value for an individual case, while horizontal lines indicate group mean values. For the *m/z* 2094 species, quantitative analysis confirms higher expression levels in MOGHE compared to TLE-HS, with the highest abundance observed in WM lesional areas from pediatric cases (N).

We first investigated two N-glycans species, Hex3HexNAc7Fuc1 and Hex3HexNAc8Fuc1, previously reported as aberrant in brain homogenates from pediatric MOGHE cases harbouring *SLC35A2* somatic mutation.^9^ The spatial distribution of these aberrant N-glycan species (defined as *m/z* 2094 and *m/z* 2297, respectively, in our dataset; Fig. 3A, B), was visualized as color-coded ion intensity maps across the tissue sections. The *m/z* 2094 signal was markedly enriched in MOGHE (Fig. 3C, F) and virtually absent in TLE-HS controls (Fig.3I; see also Supplementary Fig. 2A, B). Highest relative abundance was predominantly localized in the pediatric lesional WM, whereas adult MOGHE showed a less pronounced regional enrichment. In contrast, *m/z* 2297 displayed lower overall intensity across all cases, particularly in TLE-HS and with a milder enrichment in the lesional area in some MOGHE cases (Fig. 3D, G, L and Supplementary Fig. 2C, D). Quantitative analysis (Fig. 3N-Q) confirmed the enrichment of *m/z* 2094 in pediatric lesional WM, supporting the hypothesis of a more pronounced glycosylation alteration in pediatric than adult MOGHE cases; increased *m/z* 2094 abundance was also observed in pediatric GM despite the absence of overt histopathological abnormalities (Fig. 3N, P), consistent with the spatial molecular segmentation results.

Remarkably, within pediatric WM, the spatial distribution of the *m/z* 2094 closely mirrored the patchy hypomyelination pattern, with areas showing reduced myelin staining overlapping to regions with the highest abundance of this aberrant N-glycan (Fig. 4A-C).

**Figure 4.**
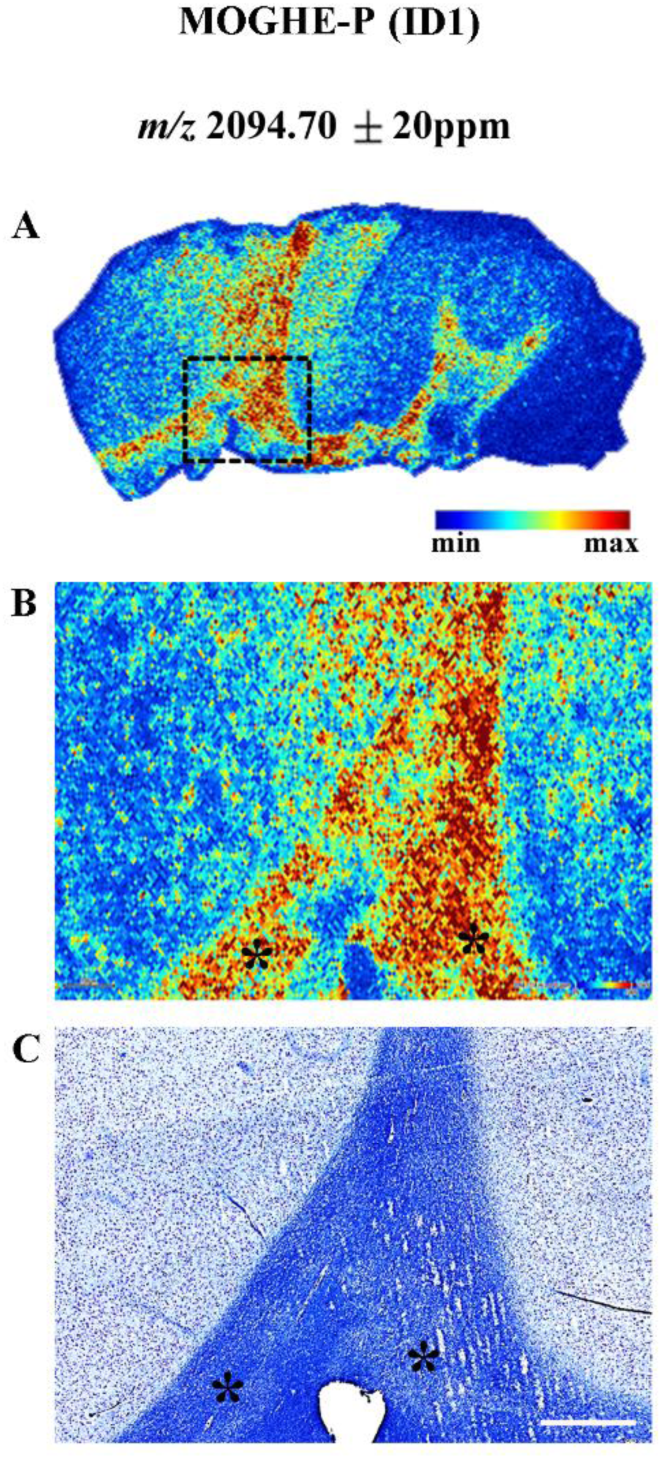
Enrichment of the aberrant *m/z* 2094 in areas of hypomyelination in pediatric MOGHE. **A**, **B**: Ion distribution map of *m/z* 2094 in a representative pediatric MOGHE case (identifier in Table 1). Box in panel **A** is shown at higher magnification in **B**. **C**: Luxol fast blu staining in an adjacent section showing the typical patchy pattern of hypomyelination (asterisks). Note the enrichment of this aberrant N-glycan (asterisks in **B**) in the regions with hypomyelination (asterisks in **C**). Scale bar: 670 micron (C)

To determine whether the spatial *m/z* 2094 distribution was part of a broader alteration of the N-glycan profile, we evaluated the abundance trajectories of all detected ions across the five tissue groups. Of the 176 ions included after matching to the reference list, 25 had *m/z* ≥3000. Most of these signals showed limited original abundance and variation across tissue groups: 18/25 (72.0%) fell below the first quartile for both mean abundance and five-group range, compared with 21/151 (13.9%) among ions below *m/z* 3000. Subsequent quantitative trend analysis was therefore restricted to the 151 ions with *m/z* <3000.

The standardized five-group abundance trajectories of the retained ions formed five recurrent profiles. Three major profiles comprised 78, 20, and 49 ions, respectively, accounting for 147 of the 151 retained signals, whereas the remaining two profiles contained only one and three ions (Supplementary Fig. 3A). In the three major profiles, both adult lesional and perilesional tissues showed closely overlapping abundances. By contrast, pediatric lesional and perilesional tissues were clearly separated in all five profiles. This finding was consistent with the molecular segmentation results, which distinguished lesional from perilesional tissue in pediatric MOGHE, but not in adult cases. The two minor trajectories contained four ions (*m/z* 2094.70, 1891.69, 1948.71, and 2297.80) all reaching their highest abundance in pediatric lesional tissue. This group included both *m/z* 2094 and *m/z* 2297, previously reported in *SLC35A2*-mutated pediatric MOGHE. Conversely, the three major trajectory profiles (147 of the 151 retained signals) were characterized either by higher abundance in pediatric perilesional tissue or by enrichment in TLE-HS.

Thus, the five trajectories could be summarized into three biologically relevant spatial profiles: pediatric lesion-enriched, pediatric perilesion-enriched, and TLE-HS-enriched. Supplementary Fig. 3B summarizes these five trajectory profiles into the three spatial profiles of interest. Fifteen ions were selected as representative of these spatial profiles by considering original signal quality, variation across tissue groups, and concordance between quantitative trajectories and spatial tissue distribution. The pediatric lesion-enriched profile comprised *m/z* 1948.71, 2094.70, 1891.69, and 2297.80. The pediatric perilesion-enriched profile included *m/z* 1752.63, 1914.68, 2207.76, and 2268.80. The TLE-HS-enriched profile comprised *m/z* 1257.42, 1955.70, 2361.86, 1996.70, 2507.91, 1850.67, and 1809.64. In the latter two profiles, adult lesional and perilesional abundances remained similar, whereas pediatric lesional tissue showed a marked decrease relative to pediatric perilesional tissue. For several of these ions, pediatric perilesional abundance was comparable to that observed in TLE-HS.

Direct evaluation of the lesion-perilesion contrasts further emphasized the age-dependent spatial differences (Supplementary Fig. 3C). In adult MOGHE, contrasts remained close to zero across the three profiles, reflecting similar abundances in lesional and perilesional tissue. In pediatric MOGHE, by contrast, lesion–perilesion differences were consistently larger and shifted away from zero.

To further examine the distribution of the 15 selected ions across GM and WM, diagnostic groups, and tissue regions, they were combined in a heatmap providing an overall view of their spatial abundance patterns (Fig. 5A).

**Figure 5.**
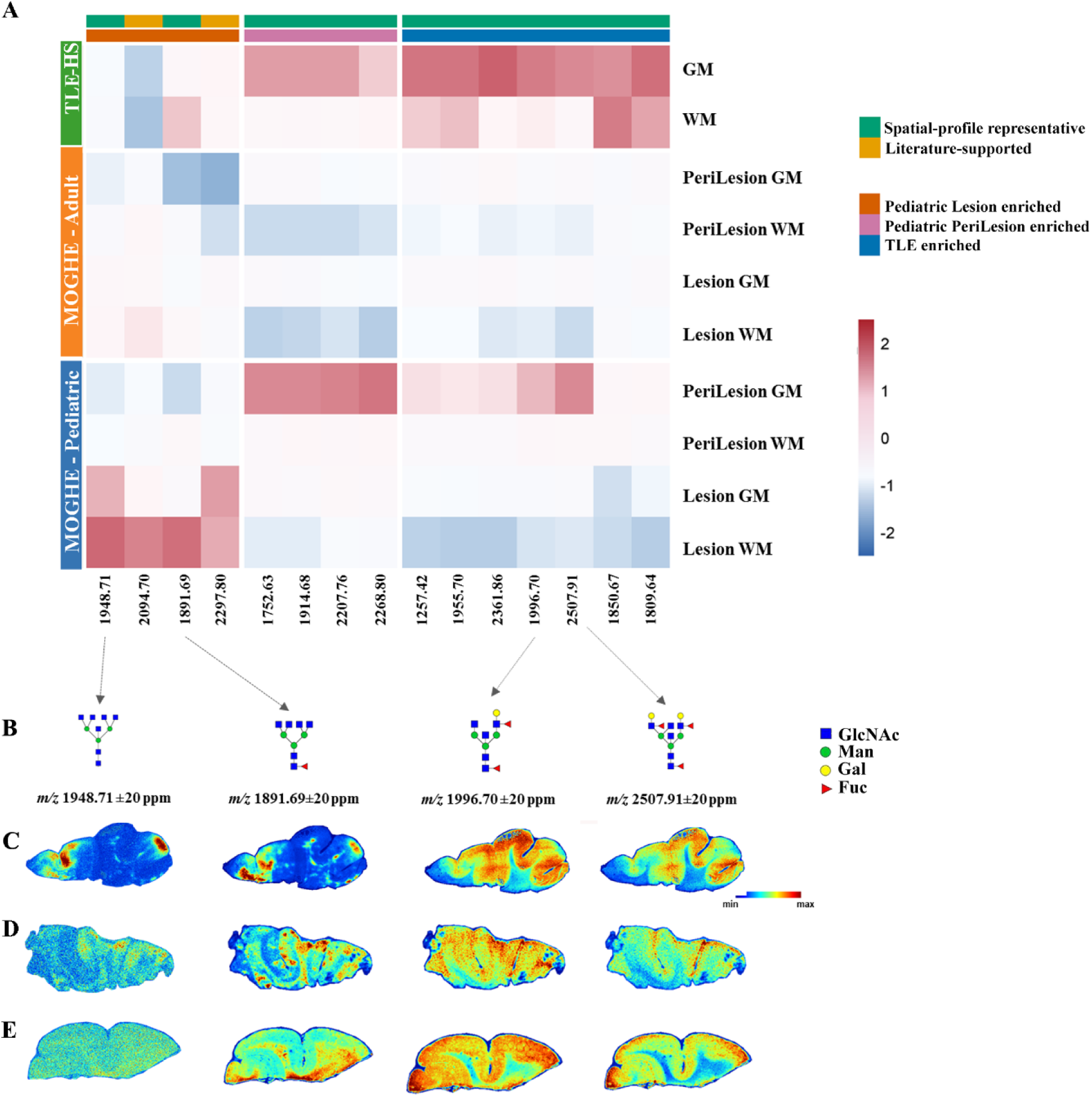
Spatially representative N-glycan abundance profiles across diagnostic groups and anatomical regions. **A**: Heatmap of the 15 representative N-glycan signals selected to summarize three spatial abundance profiles across TLE-HS, adult MOGHE perilesional and lesional tissue, and pediatric MOGHE perilesional and lesional tissue. Grey (GM) and white matter (WM) are displayed separately. Values represent z-scores obtained by standardizing the mean log₂-transformed abundance of each ion across the displayed tissue compartments. Red and blue indicate relatively high and low abundance, respectively, within each ion; colours therefore describe the spatial distribution of an individual ion and should not be used to compare absolute abundance between different ions. Columns are grouped into pediatric lesion-enriched, pediatric perilesion-enriched, and TLE-HS-enriched profiles. The upper annotations indicate the spatial profile and the selection origin of each ion. The *m/z* 2094 and *m/z* 2297 signals are indicated as literature-supported because they were previously reported in *SLC35A2*-mutated MOGHE and showed a spatial distribution consistent with the pediatric lesion-enriched profile. The remaining ions were selected as spatial-profile representatives. Row annotations indicate diagnostic macro-group, region (lesion-perilesion) and tissue compartment (WM-GM); region is not applicable to TLE-HS samples. Structures (**B**) and spatial distributions of representative glycan species in pediatric (**C**), adult MOGHE (**D**) and control TLE-HS (**E**). Notably, glycan species lacking galactose residues, including *m/z* 1948 (Hex3HexNAc7) and 1891 (Hex3dHex1HexNAc6) exhibit marked enrichment within the lesional areas of pediatric MOGHE, similarly to *m/z* 2094 described in Figure 3A. In contrast, glycan species containing one or more galactose residues, including *m/z* 1996 (Hex4dHex2HexNAc5) and *m/z* 2507 (Hex5dHex3HexNAc6), which are predominantly localized within the GM of TLE-HS control samples, show preferential enrichment in the perilesional regions of pediatric MOGHE and a more diffuse distribution pattern in adult MOGHE cases.

Boxplots of the original, non-standardized intensities confirmed the same group-specific patterns observed in the standardized trajectories (Supplementary Fig. 4). Ion maps of selected lesion-enriched and lesion-depleted species further showed that these quantitative profiles corresponded to distinct spatial distributions within the tissue (Fig. 5B-E and Supplementary Figs. 5 and 6).

Notably, in pediatric MOGHE multiantennary glycans lacking galactose residues (*m/z* 1948, 1891) were enriched in the lesional area while biantennary or multiantennary structures containing one or more galactose moieties (*m/z* 1996, 2507) were reduced (Fig. 5B-D and Supplementary Figs. 5 and 6). These findings were partially present in one adult case with high *SLC35A2* VAF. However, no correlation was found between the relative intensities of the selected ions and VAF across the MOGHE cases (Supplementary Fig. 7).

## Discussion

Glycosylation is a critical and highly dynamic post-translational modification of proteins and, through distinct biosynthetic process, of lipids. In the human brain, it plays a key role in maintaining cellular homeostasis by regulating protein folding, stability, and intracellular signalling. Glycosilation is essential during neurodevelopment, where glycan modifications of neural cell adhesion molecules contribute to neuronal differentiation and migration, and to synaptic plasticity. Through these processes, glycosylation supports the formation and refinement of neural circuits, thereby influencing interregional communication and brain-regulated physiological functions. Collectively, glycosylation acts as a fundamental molecular regulator linking neurodevelopment to the organization and function of neural networks. Despite its biological importance, brain glycosylation had been poorly characterized due to its structural heterogeneity and low abundance in neuronal tissue, and to the limited availability of robust analytical tools and comprehensive databases.^6,21^ Recent advances in mass spectrometry overcome these limitations. In particular, MALDI-MSI enables the spatially resolved analysis of N-glycan compositions within tissue sections.^13^ Using this approach, alterations in N-glycan profiles have recently been identified in brain tumours^11,12^ as well as in neurological and neurodegenerative disorders.^22–24^ These studies highlight the potential of MALDI-MSI to unravel disease-associated glycosylation patterns in the brain and to provide novel insights into the molecular mechanisms underlying neurological disorders.

Glycosylation of proteins and lipids occurs in the endoplasmic reticulum and Golgi apparatus. The *SLC35A2* gene, mutated in more than half of MOGHE patients, encodes the Golgi-localised (UDP)-galactose transporter.^25^ Analysis of microdissected cells revealed that clustered oligodendrocytes and WM heterotopic neurons are the main carriers of these variants.^5^ Accordingly, SLC35A2 protein levels localised within the Golgi apparatus are reduced in MOGHE mutated patients.^20^ Under these pathological conditions, a decreased import of UDP-galactose into the Golgi was expected, leading to impaired glycosylation of proteins.

To investigate this aspect, we analyzed the N-glycan profile in post-surgical FFPE tissue sections from six patients who underwent surgery and received a histological diagnosis of MOGHE, supported by the presence of somatic *SLC35A2* variants. Three cases of TLE-HS were included as controls. This is the first study providing *in situ* evidence of N-glycan spatially distributed alterations directly on tissue sections. Our MALDI-MSI study demonstrate: ***i)*** a distinct glycan profile that differentiates MOGHE from control TLE-HS, as well as adult MOGHE from pediatric MOGHE, ***ii)*** in pediatric cases, a distinct glycosylation pattern in, histologically-defined, lesional areas compared with perilesional regions, less evident in adult cases, and ***iii)*** the presence, particularly in pediatric cases, of a different glycan profile in the GM overlying the lesional WM, despite any evident histological alteration.

Our MOGHE cases were all characterised by a precocious age at epilepsy onset, before the age of 12, with a different history of epilepsy duration at time of surgery. The histological findings are consistent with previous reports suggesting that pediatric and adult MOGHE exhibit distinct features. In addition to oligodendroglial hyperplasia and heterotopic neurons, which are common findings, patchy areas of hypomyelination were observed exclusively in pediatric MOGHE cases. This feature has been confirmed in larger cohorts, particularly in younger patients before 8 years old, and appears to occur independent on the presence of somatic *SLC35A2* mutations in brain tissue^26,20^. Notably, this apparent paradox in younger patients, namely the coexistence of oligodendroglial hyperplasia and hypomyelination, may reflect a disruption in oligodendroglial maturation rather than a purely proliferative imbalance. It is possible that the capacity for proper myelination is at least partially restored in adulthood, although this hypothesis remains to be confirmed.

Two reports studied N-glycosylation pattern in brain MOGHE tissues. Sim et al.^9^ found that mutation-carrying samples from pediatric patients had less galactosylation, associated with truncated forms without galactose residues and aberrant N-glycan structures showing high degrees on N-acetylglucosamine (GlcNAc), compared to mutation-negative non lesional focal epilepsy and brain tumour samples. Similarly, Liu et al.^10^ found glycoproteins with aberrant N-glycosylation and reduced abundance of N-acetyllactosamine (composed of Galactose and GlcNAc), particularly in heterotopic WM neurons using a chemoenzymatic labelling.

Our N-glycan analysis performed using MALSI-MSI, has the potential to investigate the specific glycan profile within spatial context, allowing a direct correlation between the molecules and the histological features of the tissues. The method relies on-tissue application of peptide N-glycosidase F (PNGase F) to liberate N-linked glycans, which are subsequently detected to generate a glycan map directly on tissue sections.^13^

According to segmentation analysis, pediatric MOGHE cases exhibited a distinct glycosylation pattern in lesional WM, characterized by both oligodendroglial hyperplasia and hypomyelination, compared with the perilesional WM, where these histological alterations were absent. Moreover, despite the lack of overt histopathological abnormalities, the GM overlying lesional WM displayed a different glycan segmentation pattern compared to adjacent perilesional GM.

In adult MOGHE cases, lesional WM, characterized by oligodendroglial hyperplasia without hypomyelination patches, showed weaker molecular differences compared to perilesion. The quantitative trajectory analysis supported these observations: lesion-perilesion contrasts were consistently more pronounced in pediatric than in adult MOGHE tissues across the selected N-glycan profiles. Taken together, these findings suggest a more pronounced alteration of glycosylation in pediatric MOGHE compared to adult cases, and support a close relationship between myelin pathology and glycosylation patterns. The ion intensity map of *m/z* 2094, showing a particular enrichment in the patchy WM area with hypomyelination, further supports this concept.

A broader analysis showed that the observed alterations were not restricted to previously described species, such as *m/z* 2094, but involved multiple N-glycan following recurrent spatial trajectories. Representative N-glycan ions were organized into three profiles: pediatric lesion-enriched, pediatric perilesion-enriched, and TLE-HS-enriched. Adult lesional and perilesional tissues showed similar abundance levels across these profiles, whereas pediatric lesional tissue was consistently separated from the corresponding perilesional region. Thus, the main age-related difference was not a uniform increase or decrease in glycan abundance, but a stronger spatial organization of the N-glycan profile in pediatric MOGHE.

The structural composition of the three profiles was consistent with the expected consequences of impaired UDP-galactose transport. Multiantennary glycans lacking terminal galactose residues were mainly represented among pediatric lesion-enriched ions, whereas galactose-containing biantennary and multiantennary structures prevailed among pediatric perilesion- and TLE-HS-enriched ions. These findings are in agreement with previous reports of reduced galactosylation and accumulation of truncated GlcNAc-rich structures in *SLC35A2*-mutated MOGHE tissue,^9,10^ and add information on their spatial distribution within lesional, perilesional, GM, and WM compartments. These findings were consistent in all pediatric cases and were partially observed in one adult case with high percentage of VAF.

Notably, the spatially resolved nature of the method enabled the identification of a distinct glycosylation profile in MOGHE GM regions in both pediatric and adult tissues, which differed from TLE-HS controls. This represents a novel finding that warrants further investigation. The attenuated lesion-perilesion differences observed in adult MOGHE may reflect age-related changes in the molecular consequences of *SLC35A2* dysfunction or in tissue responses to chronic disease. However, the cross-sectional design and limited sample size do not allow to determine whether these findings represent either partial normalization of glycosylation, or compensatory mechanisms, or differences in disease evolution.

The functional consequences of altered N-glycosylation due to *SLC35A2* loss-of-function variants have recently been investigated in isogenic iPSC-derived neurons carrying the mutation. This *in vitro* analysis revealed disrupted early neurodevelopment, characterized by accelerated neurogenesis and altered neuronal network activity suggesting a potential link between the perturbed glycomic signature and the epileptic phenotype.^19^

In MOGHE patients, resective surgery represents a treatment option with 63% achieving a favourable surgical outcome (Engel class I) after epilepsy surgery.^27^ Interestingly, a pilot study that implemented dietary galactose supplement in a small cohort of 12 MOGHE patients with persistent seizures after surgery, demontrated clinical improvement in approximately half of the patients, particularly among those harbouring somatic mutations.^28^

The marked spatial glycosylation abnormalities observed in pediatric MOGHE provide a biological rationale for further investigation of therapies aimed at correcting galactosylation defects, including D-galactose supplementation. The substantially weaker lesion-perilesion differences observed in adult cases suggest that treatment response may vary according to age or disease duration. However, the present study was not designed to assess therapeutic efficacy, and these observations should be considered hypothesis-generating. Prospective studies integrating spatial glycomic profiles, SLC35A2 variant characteristics, age, and treatment response will be required to determine whether these molecular differences have predictive value.

## Data availability

The data that support the findings of this study are available from the corresponding author, upon reasonable request.

## Acknowledgements

The authors thank A. Cattalini for his technical support.

## Supplementary material

Supplementary material is available at Brain online’

