## Supplementary material for "Different spatial profiles of aberrant N-glycans in pediatric and adult MOGHE brain tissue": Supplemantary Material

Chiara Calabretta et al.

**Supplementary Table 1. List of genes included in the NGS panel for Focal epilepsy**

| Gene | Transcript ID | Related disease |
|---|---|---|
| MTOR | NM_004958 | FCDII (57%); HME/MEG (27%) |
| PIK3CA | NM_006218 | FCDII (1%); HME/MEG (41%) |
| AKT3 | NM_005465 | FCDII (5%); HME/MEG (14%) |
| RHEB | NM_005614 | FCDII (3%); HME/MEG (4.5%) |
| TSC1 | NM_000368 | FCDII (13.5%); HME/MEG (TSC2 |
| TSC2 | NM_000548 | 3%) |
| PTEN | NM_000314 | FCDII (1%); HME/MEG (4.5%)) |
| DEPDC5 | NM_001242896 | FCDII (13%); HME/MEG (1.5%) |
| NPRL2 | NM_006545 | FCDII (6.5%); HME/MEG (NPRL3 |
| NPRL3 | NM_001077350 | 1,5%) |
| PIK3C2B | NM_001377334 | FCDII |
| KRAS | NM_033360.4 | FCDII |
| SLC35A2 | NM_005660 | mMCD/FCD1/MOGHE |

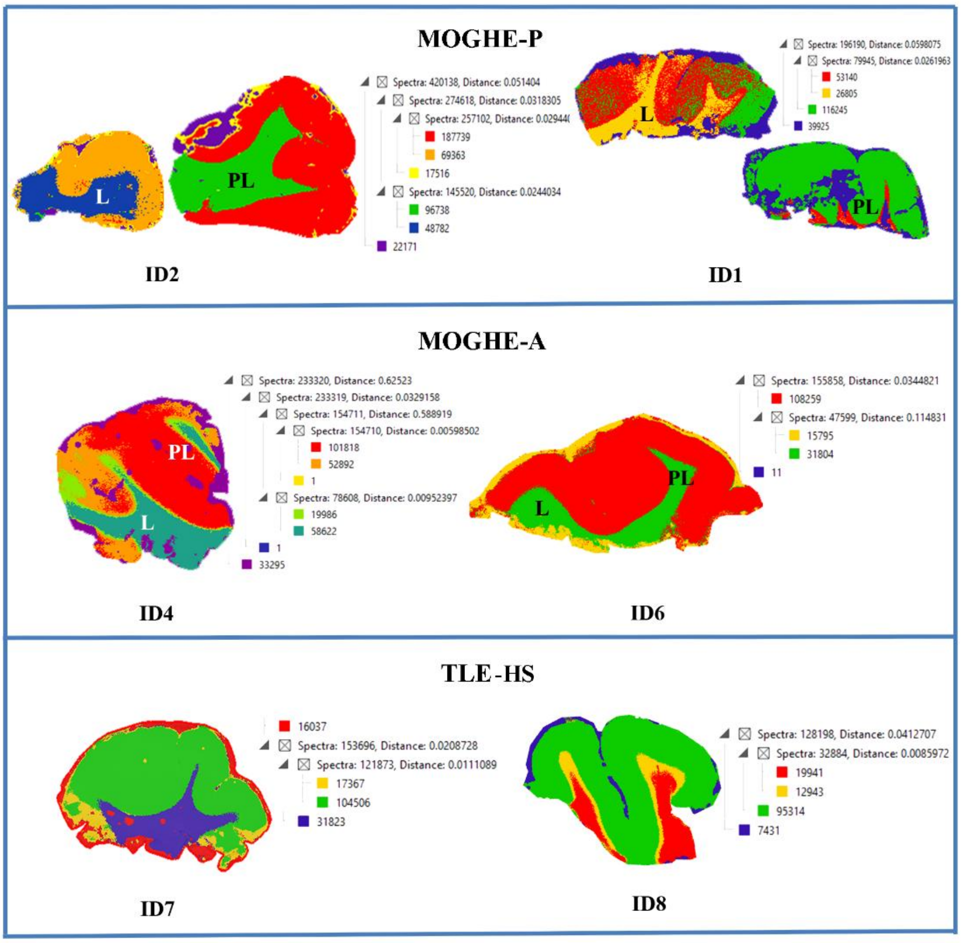


**Supplementary Figure 1**

**Supplementary Figure 1. Spatial segmentation based on N-glycan analysis: other cases.**

Spatial segmentation results in additional cases confirming the capability of the analysis to discriminate the lesional area (L) from the perilesion (PL) in pediatric cases. Mass spectra are clustered according to similarity and visualized on tissue section as color-coded segmentation maps; the corresponding dendrograms are shown on the right. Case identifiers ID correspond to those reported in Table 1.

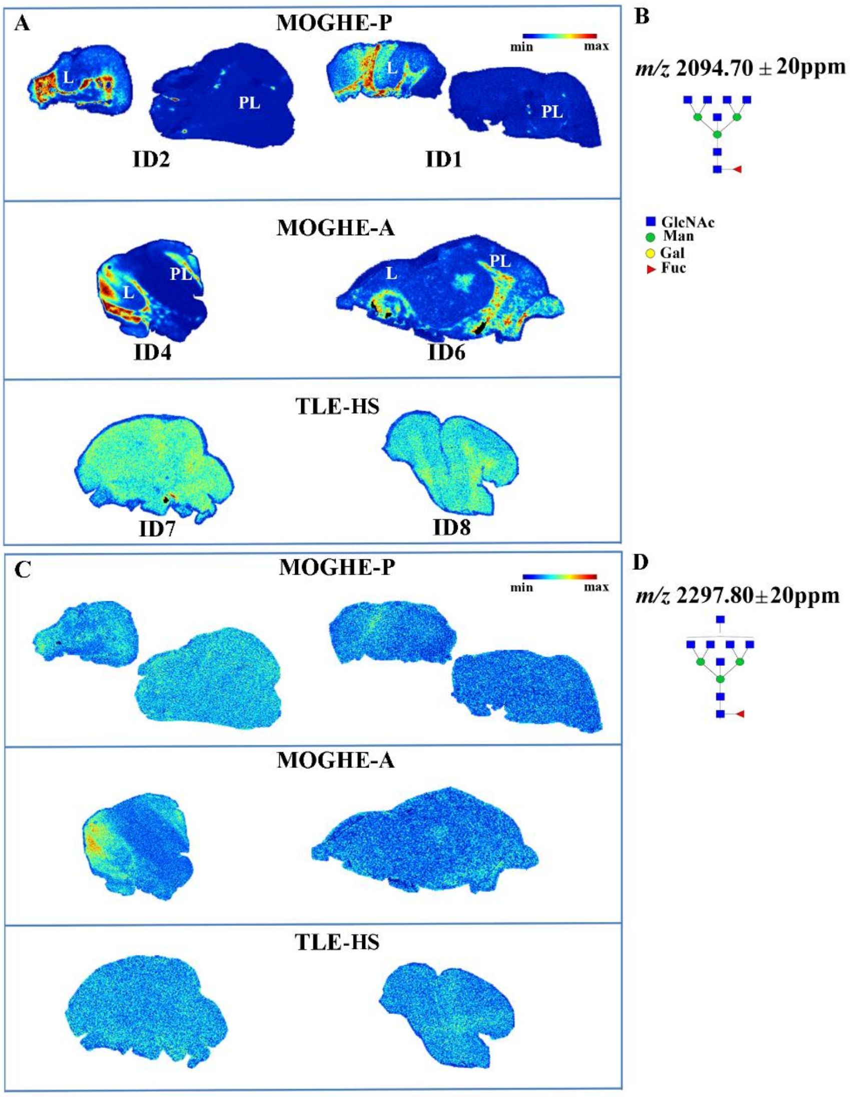


**Supplementary Figure 2**

**Supplementary Figure 2. Spatial distribution of selected, literature-supported, N-glycans in different diagnostic groups: other cases.**

**A**, **C**: Ion distribution maps of selected *m/z* species, previously reported to be altered in brain tissue from MOGHE tissue,[10] are shown in additional cases. The maps confirm the enrichment of the *m/z* 2094 (Hex3HexNAc7Fuc1, **A**) in MOGHE compared with control TLE-HS samples. In these two additional pediatric MOGHE cases, this glycan is predominantly enriched in the lesional area (L) relative to the perilesion (PL), whereas this distribution pattern is less consistent in adult cases. In contrast, *m/z* 2297 (Hex3HexNAc8Fuc1, **C**) is detected at low abundance across all cases, particularly in TLE-HS samples and with a milder enrichment in the lesional area in some MOGHE cases. Case identifiers ID correspond to those reported in Table 1. The corresponding glycan structures and legend are represented in **B** and **D**, respectively. Relative intensity quantification across all cases is represented in Fig. 3.

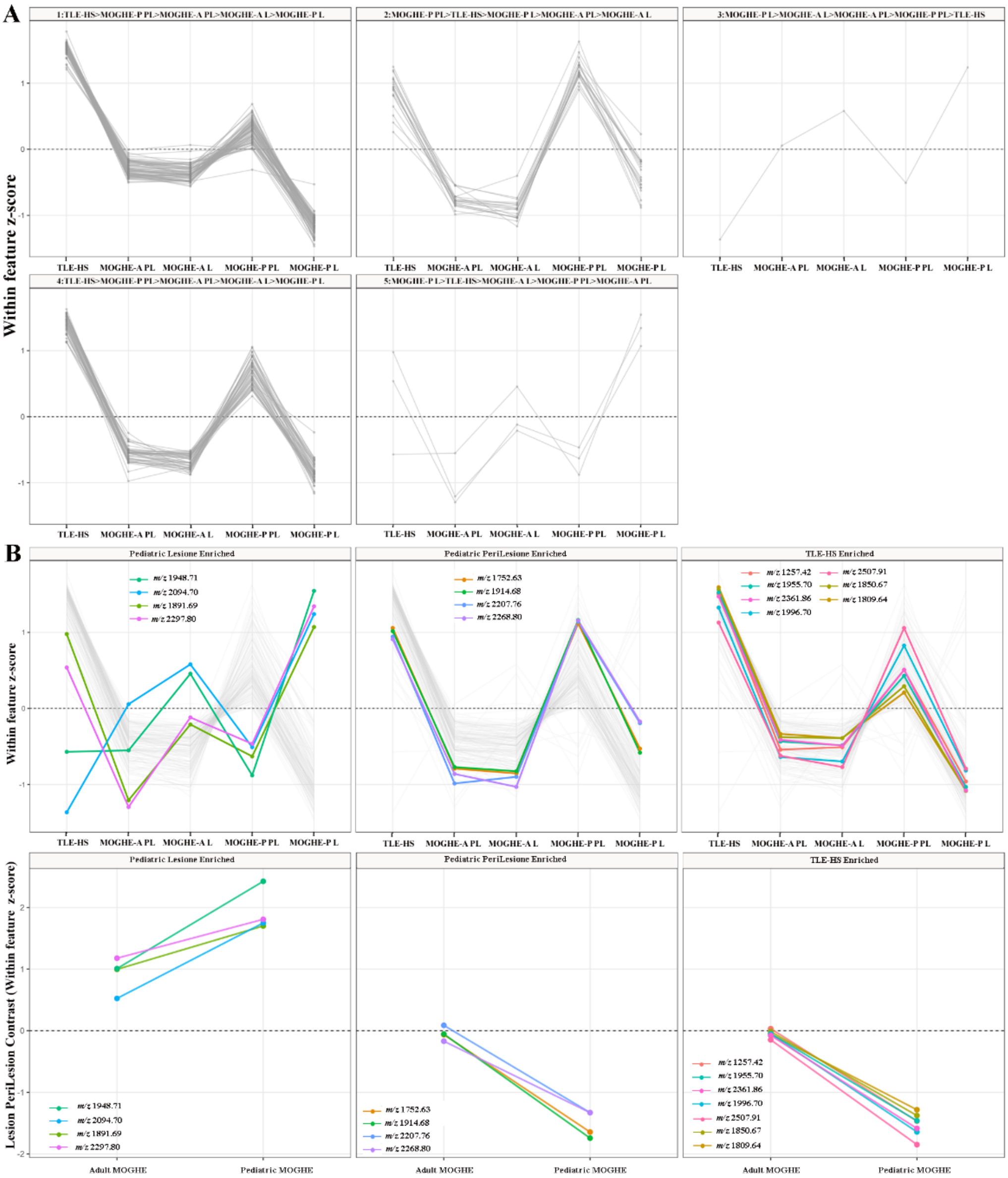


**Supplementary figure 3**

**Supplementary Figure 3. Global and selected spatial N-glycan abundance trajectories.**

**A**: Global abundance trajectories of all evaluated N-glycan signals with *m/z* <3000. Ions were grouped into five clusters according to the similarity of their standardized abundance trajectories across TLE-HS, adult MOGHE perilesional tissue, adult MOGHE lesional tissue, pediatric MOGHE perilesional tissue, and pediatric MOGHE lesional tissue. Each grey line represents one ion, and facet labels report the complete ordering of the five group means within each cluster.

**B:** Abundance trajectories of the 15 ions included in the final heatmap, grouped into the three spatial profiles: pediatric lesion-enriched, pediatric perilesion-enriched, and TLE-HS-enriched. Selected ions are shown as coloured lines, whereas all other eligible ions with *m/z* <3000 are shown in grey to place the selected profiles within the complete dataset.

**C:** Lesion–perilesion contrasts for the selected ions in adult and pediatric MOGHE. Contrasts were calculated as the difference between mean standardized abundance in lesional and perilesional tissue. Positive values indicate lesion enrichment, whereas negative values indicate perilesion enrichment. Each line represents one selected ion. Adult contrasts remained close to zero across the three profiles, whereas pediatric contrasts showed a consistent displacement from zero.

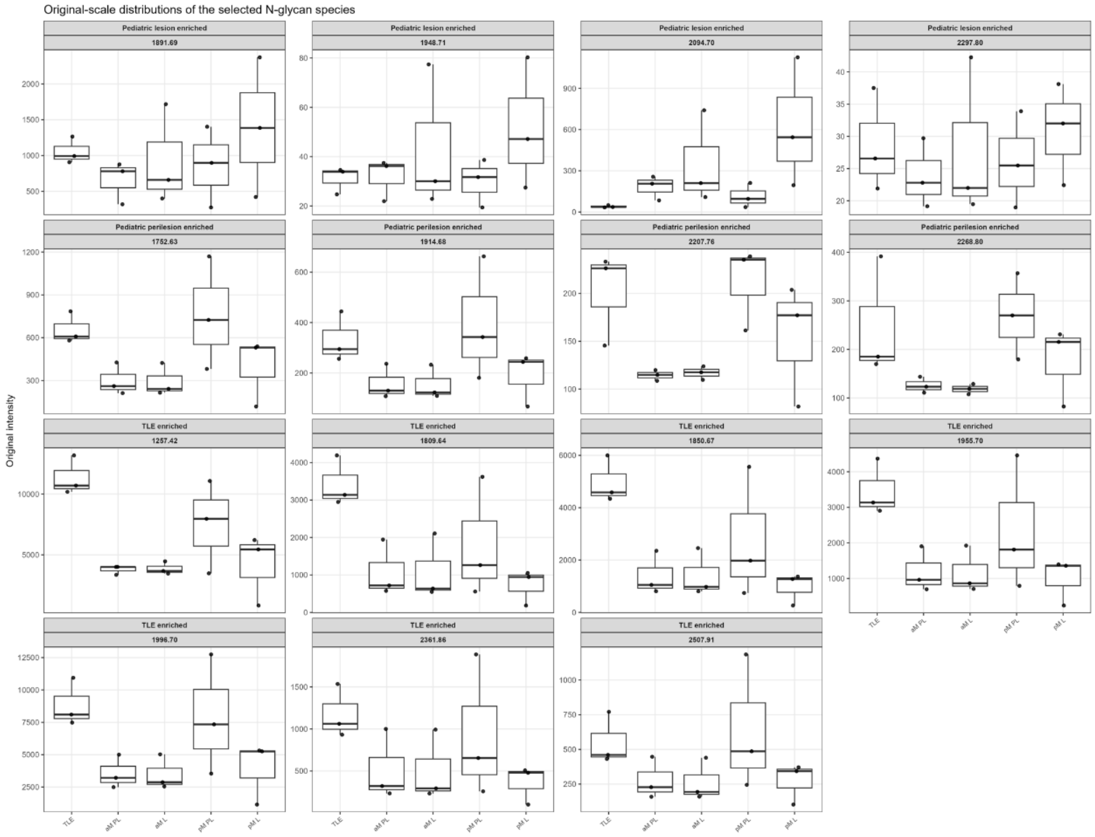


**Supplementary figure 4**

**Supplementary Figure 4. Boxplots of the original, non-standardized intensities.**

Original, non-standardized MALDI-MSI intensities of the selected ions across the five tissue groups. Points represent patient-level abundance values. These distributions allow the spatial trajectories and lesion–perilesion differences identified after within-feature standardization to be evaluated on the original intensity scale. Abbreviations: aM, adult MOGHE; pM, pediatric MOGHE; L, lesional; PL, perilesional; TLE-HS, temporal lobe epilepsy with hippocampal sclerosis.

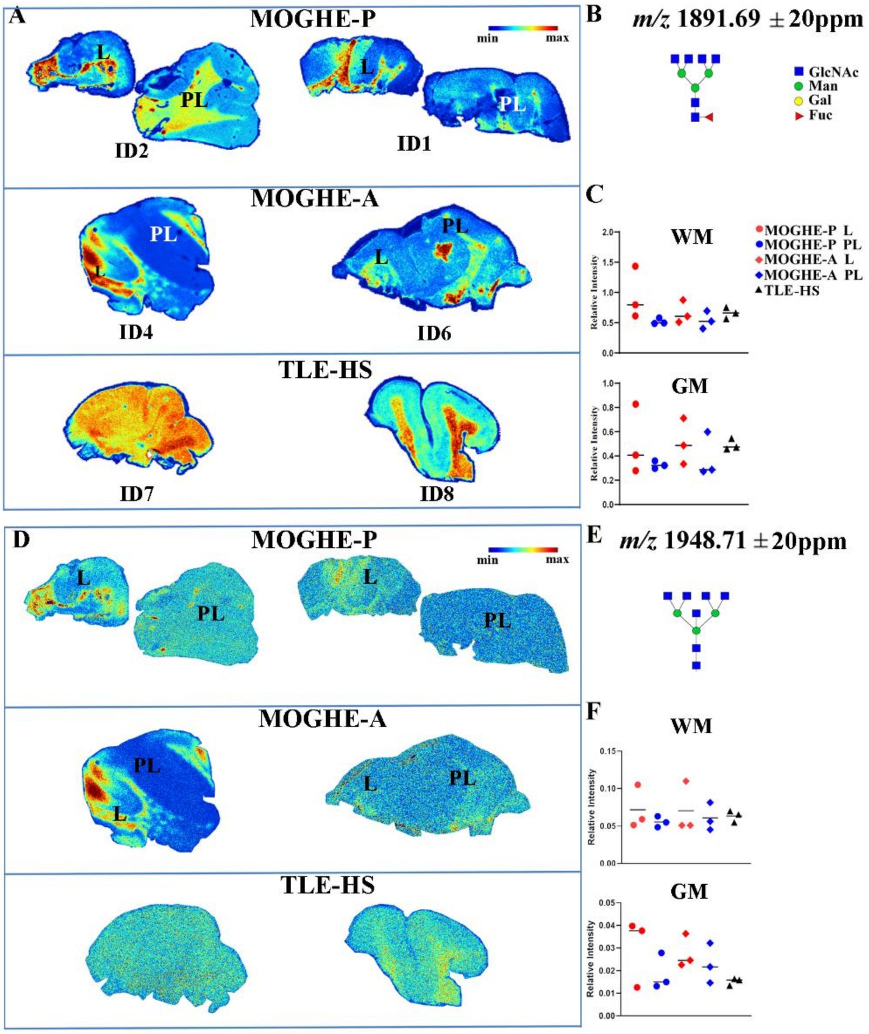


**Supplementary Figure 5**

**Supplementary Figure 5. Ion distribution maps of specific *m/z* glycan species.**

Ion distribution maps of *m/z* 1891 (Hex3dHex1HexNAc6, **A**) and *m/z* 1948 (Hex3HexNAc7 **D**), glycan species lacking galactose residues, in additional cases. **C**, **F**: Relative intensity measurements of these two *m/z* species across different ROIs in all cases. Both maps and quantifications confirm the enrichment within the lesional areas of pediatric MOGHE, particularly for *m/z* 1891, whereas this pattern is less consistent in adult cases. Case identifiers ID correspond to those reported in Table 1. The corresponding glycan structures and legend are shown in **B** and **E**, respectively. In the plots (C, F), each dot represents the mean intensity of the indicated *m/z* species within a given ROI, while the horizontal line indicates the group mean.

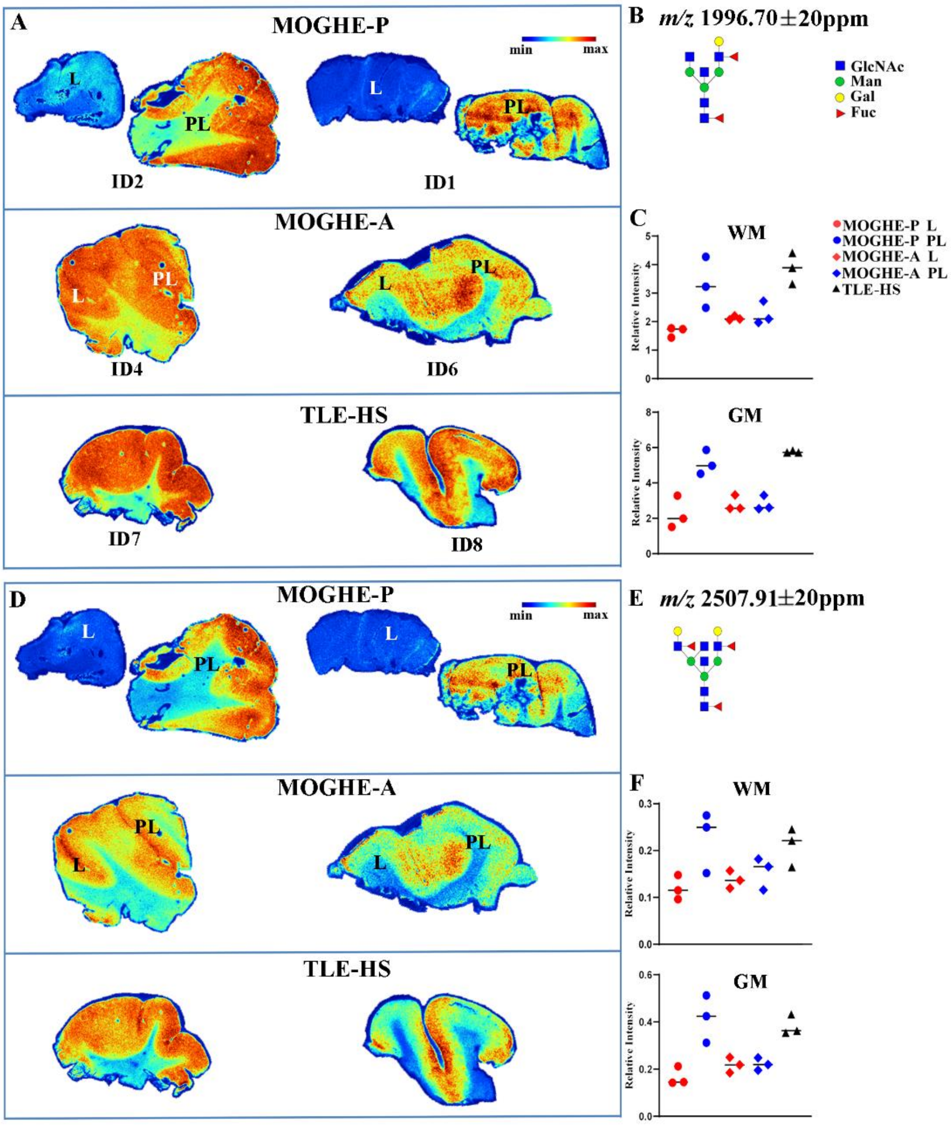


Supplementary Figure 6

**Supplementary Figure 6. Ion distribution maps of specific *m/z* glycan species.**

**A**, **D**: Ion distribution maps of *m/z* 1996 (Hex4dHex2HexNAc5; **A**) and *m/z* 2507 (Hex5dHex3HexNAc6; **D**), both containing galactose moieties, are shown in additional cases. **C**, **F**: Relative intensity quantification of these two *m/z* species across different ROIs in all cases. Both the ion distribution maps and quantitative analyses confirm decreased levels in lesion compared to perilesional regions of pediatric MOGHE and to TLE-HS samples, especially in GM. Adult MOGHE tissues exhibit intermediate abundance levels. Case identifiers ID correspond to those reported in Table 1. The corresponding glycan structures and legend are shown in **B** and **E**, respectively. In the plots (C, F), each dot represents the mean intensity of the indicated m/z species within a given ROI, while the horizontal line indicates the group mean.

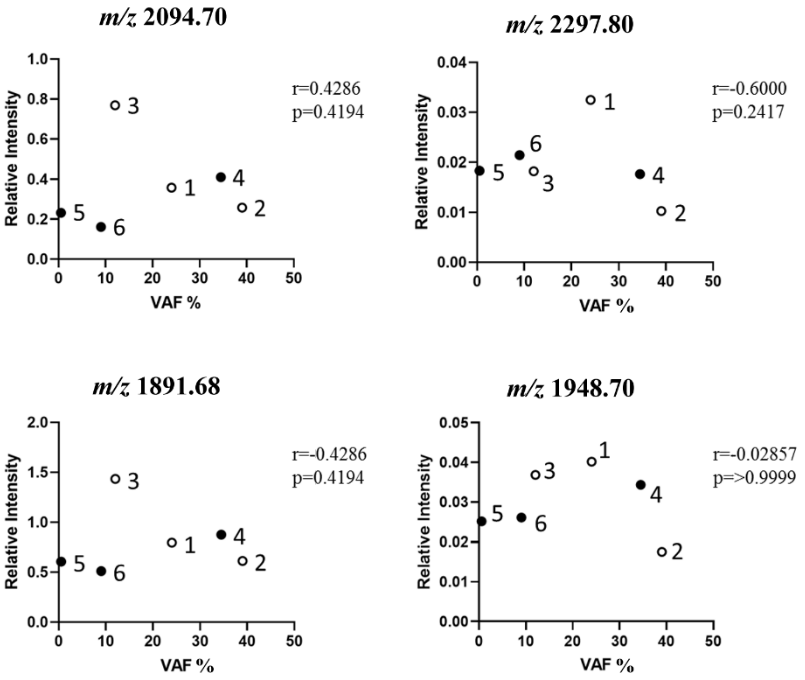


### Supplementary Figure 7

**Supplementary Figure 7. Spearman analysis of correlation between ion intensity and VAF of *SLC35A2*.**

Graphs showing variant allele frequency (VAF) plotted against the relative ion intensity in the lesional white matter area were generated for *m/z* 2094 and *m/z* 2297, previously reported to be altered in MOGHE (Sim et al., 2018), as well as for *m/z* 1891 and *m/z* 1948, identified as discriminating features in the present study, across all MOGHE samples. Spearman's correlation coefficients (r) and associated p values indicate no significant correlation between VAF and relative ion intensity for any of the analyzed *m/z* species. Open circles represent pediatric MOGHE cases, whereas filled circles represent adult MOGHE cases. Case identifier ID numbers on plots correspond to those reported in Table 1.

**Supplementary Table 2. Signal-quality assessment, global trajectory clusters, and quantitative characteristics of the N-glycan ions selected to represent the three spatial abundance profiles.** The file contains four worksheets. *Selection_notes* summarizes the rationale for signal-quality assessment, trajectory clustering, and selection of the representative ions. *Signal_QC* reports the number and proportion of ions classified as having both low original abundance and limited biological dynamic range, separately for signals with *m/z* <3000 and *m/z* ≥3000, together with the first-quartile thresholds used for this assessment. *Trajectory_clusters* describes the five recurrent abundance trajectories identified among ions with *m/z* <3000, reporting the number of ions assigned to each cluster and the mean standardized abundance across TLE-HS, adult MOGHE perilesional and lesional tissue, and pediatric MOGHE perilesional and lesional tissue. *Selected_15_m/z* reports the displayed and measured *m/z* values, assigned spatial profile, trajectory cluster, selection origin, original signal-quality metrics, detection rate, mean abundance in each of the five tissue groups, adult and pediatric lesion–perilesion contrasts, epsilon-squared effect size, Kruskal–Wallis p-value, and Benjamini–Hochberg-adjusted p-value for the 15 ions included in the final heatmap.

Lesion–perilesion contrasts were calculated as lesional minus perilesional standardized abundance; positive values indicate lesion enrichment, whereas negative values indicate perilesion enrichment. Statistical comparisons were performed across the three independent macro-groups—TLE-HS, adult MOGHE, and pediatric MOGHE—after patient-level aggregation and should be interpreted as exploratory because of the limited cohort size.
