## Supplementary Table 2 for "Different spatial profiles of aberrant N-glycans in pediatric and adult MOGHE brain tissue"

**Selection notes**

| Item | Description |
| --- | --- |
| Purpose | This file reports the signal-quality assessment, the five global trajectory clusters, and the characteristics of the 15 representative ions shown in the final heatmap. |
| Signal-quality assessment | Original mean abundance and the range across the five tissue groups were evaluated before within-feature standardization. Ions at or above m/z 3000 were excluded because this mass range was predominantly represented by signals below the first quartile of both measures. |
| Trajectory clustering | After restriction to ions below m/z 3000, five recurrent trajectory groups were represented in the retained dataset. |
| Final spatial profiles | Fifteen ions were selected as spatially representative of three biological profiles: pediatric lesion-enriched, pediatric perilesion-enriched, and TLE-HS-enriched. |
| Statistical analysis | Kruskal-Wallis comparisons were performed across the three independent macrogroups after patient-level aggregation. Epsilon squared describes the magnitude of the group effect; p-values were adjusted using the Benjamini-Hochberg method. |
| Literature-supported ions | m/z 2094 and m/z 2297 were retained because they had been previously reported in SLC35A2-mutated MOGHE and their observed spatial distributions were compatible with the pediatric lesion-enriched profile. |
| Interpretation of contrasts | Adult and pediatric L-PL contrasts are calculated from the five-group standardized trajectories. Positive values indicate lesion enrichment; negative values indicate perilesion enrichment. |

**Signal QC**

| Mass_group | N_ions | N_low_abundance_and_range | Percentage_low_abundance_and_range | Median_mean_original_intensity | Median_five_group_range | Mean_abundance_Q1_cutoff | Five_group_range_Q1_cutoff |
| --- | --- | --- | --- | --- | --- | --- | --- |
| m/z < 3000 | 151 | 21.0 | 13.907 | 126.517 | 150.484 | 51.640 | 32.556 |
| m/z ≥ 3000 | 25 | 18.0 | 72.000 | 39.291 | 23.820 | 51.640 | 32.556 |

**Trajectory clusters**

| Band_number | N_ions | Complete_order | Mean_z_TLE | Mean_z_aMPL | Mean_z_aML | Mean_z_pMPL | Mean_z_pML |
|---|---|---|---|---|---|---|---|
| 1 | 78 | TLE > pM PL > aM PL > aM L > pM L | 1.524 | -0.298 | -0.366 | 0.280 | -1.139 |
| 2 | 20 | pM PL > TLE > pM L > aM PL > aM L | 0.884 | -0.768 | -0.906 | 1.199 | -0.410 |
| 4 | 1 | pM L > aM L > aM PL > pM PL > TLE | -1.366 | 0.056 | 0.579 | -0.509 | 1.240 |
| 5 | 49 | TLE > pM PL > aM PL > aM L > pM L | 1.407 | -0.567 | -0.675 | 0.668 | -0.833 |
| 6 | 3 | pM L > TLE > aM L > pM PL > aM PL | 0.315 | -1.018 | 0.042 | -0.658 | 1.319 |

### Selected 15 m/z

| Figure m/z | Measured m/z | Spatial_profile | Band_number | Complete_order | Selection_basis | Literature_supported | Mean_original_abundance |
|---|---|---|---|---|---|---|---|
| 1948.71 | 1948.71 | Pediatric lesion enriched | 6 | pM L > TLE > aM L > pM PL > aM PL | Spatial-profile representative | No | 37.581 |
| 2094.70 | 2094.70 | Pediatric lesion enriched | 4 | pM L > aM L > aM PL > pM PL > TLE | Literature-supported | Yes | 262.136 |
| 1891.69 | 1891.69 | Pediatric lesion enriched | 6 | pM L > TLE > aM L > pM PL > aM PL | Spatial-profile representative | No | 977.629 |
| 2297.80 | 2297.80 | Pediatric lesion enriched | 6 | pM L > TLE > aM L > pM PL > aM PL | Literature-supported | Yes | 27.485 |
| 1752.63 | 1752.63 | Pediatric perilesion enriched | 2 | pM PL > TLE > pM L > aM PL > aM L | Spatial-profile representative | No | 481.058 |
| 1914.68 | 1914.68 | Pediatric perilesion enriched | 2 | pM PL > TLE > pM L > aM PL > aM L | Spatial-profile representative | No | 246.061 |
| 2207.76 | 2207.76 | Pediatric perilesion enriched | 2 | pM PL > TLE > pM L > aM PL > aM L | Spatial-profile representative | No | 159.837 |
| 2268.80 | 2268.80 | Pediatric perilesion enriched | 2 | pM PL > TLE > pM L > aM PL > aM L | Spatial-profile representative | No | 187.676 |
| 1257.42 | 1257.42 | TLE enriched | 5 | TLE > pM PL > aM PL > aM L > pM L | Spatial-profile representative | No | 6133.033 |
| 1955.70 | 1955.70 | TLE enriched | 1 | TLE > pM PL > aM PL > aM L > pM L | Spatial-profile representative | No | 1835.074 |
| 2361.86 | 2361.86 | TLE enriched | 1 | TLE > pM PL > aM PL > aM L > pM L | Spatial-profile representative | No | 698.968 |
| 1996.70 | 1996.69 | TLE enriched | 5 | TLE > pM PL > aM PL > aM L > pM L | Spatial-profile representative | No | 5533.830 |
| 2507.91 | 2507.91 | TLE enriched | 5 | TLE > pM PL > aM PL > aM L > pM L | Spatial-profile representative | No | 401.122 |
| 1850.67 | 1850.67 | TLE enriched | 1 | TLE > pM PL > aM PL > aM L > pM L | Spatial-profile representative | No | 2302.757 |
| 1809.64 | 1809.64 | TLE enriched | 1 | TLE > pM PL > aM PL > aM L > pM L | Spatial-profile representative | No | 1628.366 |

| Five_group_range | Detection_percent | TLE mean | Adult PL mean | Adult L mean | Pediatric PL mean | Pediatric L mean | Adult_L_minus_PL | Pediatric_L_minus_PL | Epsilon_squared | Kruskal_Wallis_p_value | FDR |
|---|---|---|---|---|---|---|---|---|---|---|---|
| 21.699 | 100.0 | 31.053 | 31.859 | 43.443 | 29.926 | 51.625 | 1.008 | 2.425 | 0.033 | 0.875 | 0.880 |
| 582.576 | 100.0 | 39.380 | 182.366 | 353.032 | 113.948 | 621.956 | 0.524 | 1.749 | 0.678 | 0.066 | 0.234 |

| | | | | | | | | | | | |
|---|---|---|---|---|---|---|---|---|---|---|---|
| 733.703 | 100.0 | 1054.305 | 658.360 | 925.448 | 857.968 | 1392.063 | 0.996 | 1.701 | 0.144 | 0.561 | 0.599 |
| 6.964 | 100.0 | 28.661 | 23.887 | 27.911 | 26.116 | 30.851 | 1.177 | 1.808 | 0.044 | 0.837 | 0.847 |
| 465.151 | 100.0 | 657.928 | 299.810 | 293.271 | 758.422 | 395.861 | -0.063 | -1.645 | 0.433 | 0.177 | 0.324 |
| 240.204 | 100.0 | 331.719 | 158.317 | 155.087 | 395.290 | 189.893 | -0.055 | -1.744 | 0.478 | 0.148 | 0.324 |
| 97.380 | 100.0 | 201.700 | 114.431 | 116.986 | 211.811 | 154.256 | 0.087 | -1.330 | 0.544 | 0.113 | 0.273 |
| 150.484 | 100.0 | 248.890 | 125.956 | 118.381 | 268.865 | 176.286 | -0.170 | -1.330 | 0.544 | 0.113 | 0.273 |
| 7577.277 | 100.0 | 11358.898 | 3781.622 | 3859.303 | 7505.919 | 4159.425 | 0.033 | -1.467 | 0.700 | 0.061 | 0.234 |
| 2472.047 | 100.0 | 3469.351 | 1188.526 | 1165.205 | 2354.984 | 997.304 | -0.050 | -1.462 | 0.678 | 0.066 | 0.234 |
| 814.450 | 100.0 | 1176.371 | 518.014 | 505.664 | 932.871 | 361.921 | -0.077 | -1.587 | 0.411 | 0.193 | 0.324 |
| 5357.409 | 100.0 | 8838.855 | 3567.659 | 3481.446 | 7875.317 | 3905.871 | -0.059 | -1.642 | 0.478 | 0.148 | 0.324 |
| 373.948 | 100.0 | 553.756 | 277.639 | 264.212 | 638.160 | 271.843 | -0.148 | -1.849 | 0.344 | 0.252 | 0.400 |
| 4006.689 | 100.0 | 4974.112 | 1402.346 | 1410.995 | 2758.908 | 967.423 | -0.015 | -1.377 | 0.678 | 0.066 | 0.234 |
| 2698.415 | 100.0 | 3425.018 | 1079.724 | 1097.774 | 1812.708 | 726.603 | -0.054 | -1.283 | 0.678 | 0.066 | 0.234 |
